# Immunogen selection and prior immunity shape antibody breadth following immunisation with avian H5 hemagglutinin

**DOI:** 10.64898/2026.09.01.748495

**Authors:** Yee-Chen Liu, Andrew Kelly, Robyn Esterbauer, Wen Shi Lee, Stephen J. Kent, Marios Koutsakos, Adam K. Wheatley

## Abstract

Avian influenza A viruses pose a persistent zoonotic threat to humans owing to their expanding host range and high case fatality rates. In particular, viruses from the 2.3.4.4b clade of the H5 subtype have now been detected in over 60 mammalian species, raising serious pandemic concerns. Understanding immune recognition of the H5 hemagglutinin (HA) is therefore critical for effective vaccine design and pandemic preparedness. To understand the breadth of cross-recognition induced by different H5 strains, we selected genetically diverse H5 human isolates from 2003-2023 and assessed neutralising antibody responses elicited by adjuvanted recombinant HA protein-based vaccines in C57BL/6 mice. Neutralisation activity of sera was determined against seven H5 HA variants using pseudotyped viruses and a PR8-reassortant virus in micro-neutralisation assays. Our results showed a wide variety of cross-strain neutralisation across H5 HA antigen variants. The conventional vaccine strain A/Indonesia/05/2005 displayed narrow activity against emerging clade 2.3.4.4b viruses, whereas ancestral variants exhibited cross-neutralisation profiles showing a diversity of breath but with limited potency. Polyvalent H5 HA formulations and nanoparticle-displayed H5 HA platforms substantially broadened cross-neutralisation against diverse H5 strains. To examine the impact of pre-existing immunity on H5 vaccine immunogenicity in mouse models, mice were primed with either seasonal influenza infection or quadrivalent influenza vaccine (QIV) prior to H5 HA immunisation. QIV pre-vaccination, but not prior influenza infection, enhanced subsequent neutralizing responses towards A/Fujian-Sanyuan/21099/2017 (clade 2.3.4.4b) H5. Collectively, our results demonstrate that immunogen selection and prior immunity shape antibody breadth following immunisation with avian A(H5) hemagglutinin.

**Importance:** Highly pathogenic avian influenza H5 viruses continue to spread across an unprecedented range of mammalian hosts, heightening the risk of a human pandemic. Current vaccine approaches for H5 rely on frequent recommendations of candidate vaccine viruses to match emerging H5 strains. Developing broadly protective H5 vaccines is thus a priority as part of pandemic preparedness. This study demonstrates a considerable variability in the potential of H5 vaccine antigens to induce neutralisation breadth, as well as the potential for multivalent vaccines and ferritin nanoparticle-based strategies to robustly augment immunity across antigenic variants. Furthermore, prior immunity established by seasonal QIV impacts the immunogenicity of subsequent 2.3.4.4b HA vaccination against H5 diversity. These findings provide critical insights to guide the rational design of broadly reactive H5 vaccines and inform pre-pandemic preparedness strategies.

## Introduction

Avian influenza A viruses (IAV) pose a persistent zoonotic threat to human health due to their broad host range, recurrent outbreaks, and high potential for morbidity and mortality (1). This is particularly true for high pathogenicity avian influenza viruses (HPAIV) of the A(H5) Goose/Guandong (Gs/Gd) lineage that emerged in 1997. The continuous circulation of Gs/Gd H5 viruses over the years has resulted in considerable diversification into multiple genetic clades. Since 2021, spillovers involving clade 2.3.4.4b H5Nx viruses have been documented in more than 60 mammalian species (2, 3), raising concerns about the risk of viral adaptation and the potential for disseminated mammal-to-mammal transmission. In 2024, the spread of clade 2.3.4.4b H5 viruses into U.S. dairy cattle led to substantial economic losses and sporadic human infections (4), which to date have remained associated with low pathogenicity in humans. In parallel, human infections caused by H5 HPAIV from clade 2.3.2.1e were reported in Cambodia between 2023 and 2024, linked to the emergence of a novel reassortant genotype (5). The significant and ongoing risk of avian influenza necessitates a better understanding of immune recognition of avian influenza viruses, and strategies to maximise the effectiveness of vaccines or treatments for responding to future pandemics.

Influenza vaccines typically target the haemagglutinin (HA) as the major surface glycoprotein that mediates receptor binding and membrane fusion during viral entry. Immune pressure from antibodies drives antigenic diversification of the HA, mediated by mutations in residues surrounding the receptor-binding site (RBS) of both seasonal and avian influenza viruses (6–8). The resulting antigenic diversity of A(H5) viruses complicates pandemic preparedness and the stockpiling of broadly effective vaccines. H5 vaccine development has largely followed the seasonal influenza vaccine paradigm, guided by continuous global surveillance and antigenic characterisation coordinated through the WHO Global Influenza Surveillance and Response System, with frequent recommendations of candidate vaccine virus (CVV) strains to optimise matching with circulating strains (9–11). At least 20 H5 vaccine products are licensed across America, Asia, Australia, and Europe and are predominantly inactivated, egg-based formulations requiring adjuvants and two-dose regimens (12). The antibody response induced has typically been assessed against a limited number of historical H5 isolates, primarily from clades 1 and 2.1 or 2.3.4. It is therefore not clear to what extent such vaccines provide coverage against the full breadth of H5 antigenic diversity (13).

Given the impracticalities of continuous CVV re-selection, alternative strategies to induce H5 subtype-wide immunity are being explored. Recent advances include high-resolution antigenic mapping to identify “central” vaccine strains (14), next-generation platforms such as self-replicating mRNA vaccines capable of rapid, flexible, and multivalent antigen design (15, 16), heterologous prime-boost strategies (17, 18), and computationally optimised broadly reactive antigens (COBRA) (19, 20). These strategies highlight both the opportunities and the continuing need for vaccine approaches capable of overcoming H5 antigenic diversity and achieving broad, durable protection against this subtype of pandemic potential.

In this study, we explored the impact of immunogen selection and pre-existing immunity upon the breadth of antibody responses elicited against H5Nx viruses in a mouse model. Our findings suggest (i) considerable variability in the potential of H5 vaccine antigens to induce neutralisation breadth, (ii) the potential for multivalent vaccines and ferritin nanoparticle-based strategies to robustly augment immunity across antigenic variants; and (iii) a role for prior immunity established by seasonal QIV to impact the breadth and potency of cross-neutralisation activity in response to immunisation with 2.3.4.4b HA.

## Results

### Different H5 strains elicit heterogeneous cross-strain neutralisation

We firstly wanted to understand the potential for cross-strain neutralisation that can be induced after immunisation with different H5 strains. Eight H5Nx viral isolates from human zoonotic infections were selected that span the genetic and antigenic diversity of the H5 Gs/Gd lineage: A/Indonesia/5/2005 (Indo/05; clade 2.1.3.2), A/Anhui/1/2005 (An/05; clade 2.3.4), A/Egypt/N0423/2011 (Egy/11; clade 2.2.1), A/Cambodia/X1030304/2013 (Cam/13; clade 1.1.2), A/Cambodia/2302009/2023 (Cam/23; clade 2.3.2.1c), A/Sichuan/26221/2014 (Sic/14; clade 2.3.4.4a), A/Fujian-Sanyuan/21099/2017 (Fuj/17; clade 2.3.4.4b) and A/Hubei/29578/2016 (Hub/16; clade 2.3.4.4d) (Fig. 1). Recombinant, trimeric soluble HA proteins were generated for each strain as described (21) and used to vaccinate C57BL/6 mice using a two-dose prime-boost regimen of HA formulated with AddaVax, an MF-59-like adjuvant. An HA from B/Lee/40 served as a negative comparator. Serum neutralisation activity against an H5 clade 2.3.4.4b virus was measured in a micro-neutralisation assay using a recombinant IAV comprising HA and NA genes from A/Astrakhan/3212/2020 (Ast/20) and internal genes from H1 PR8. As such PR8 recombinant viruses were not available for other H5 strains, we instead utilised lentivirus-based pseudoviruses displaying the HAs from Indo/05, An/05, Egy/11, Cam/13, Cam/23, and Hub/16 to measure neutralisation (Fig. 2a; Supp. Fig. 1; Supp. Fig. 2). The sera from each H5 antigen group were titrated against all the viruses.

**Figure 1.**
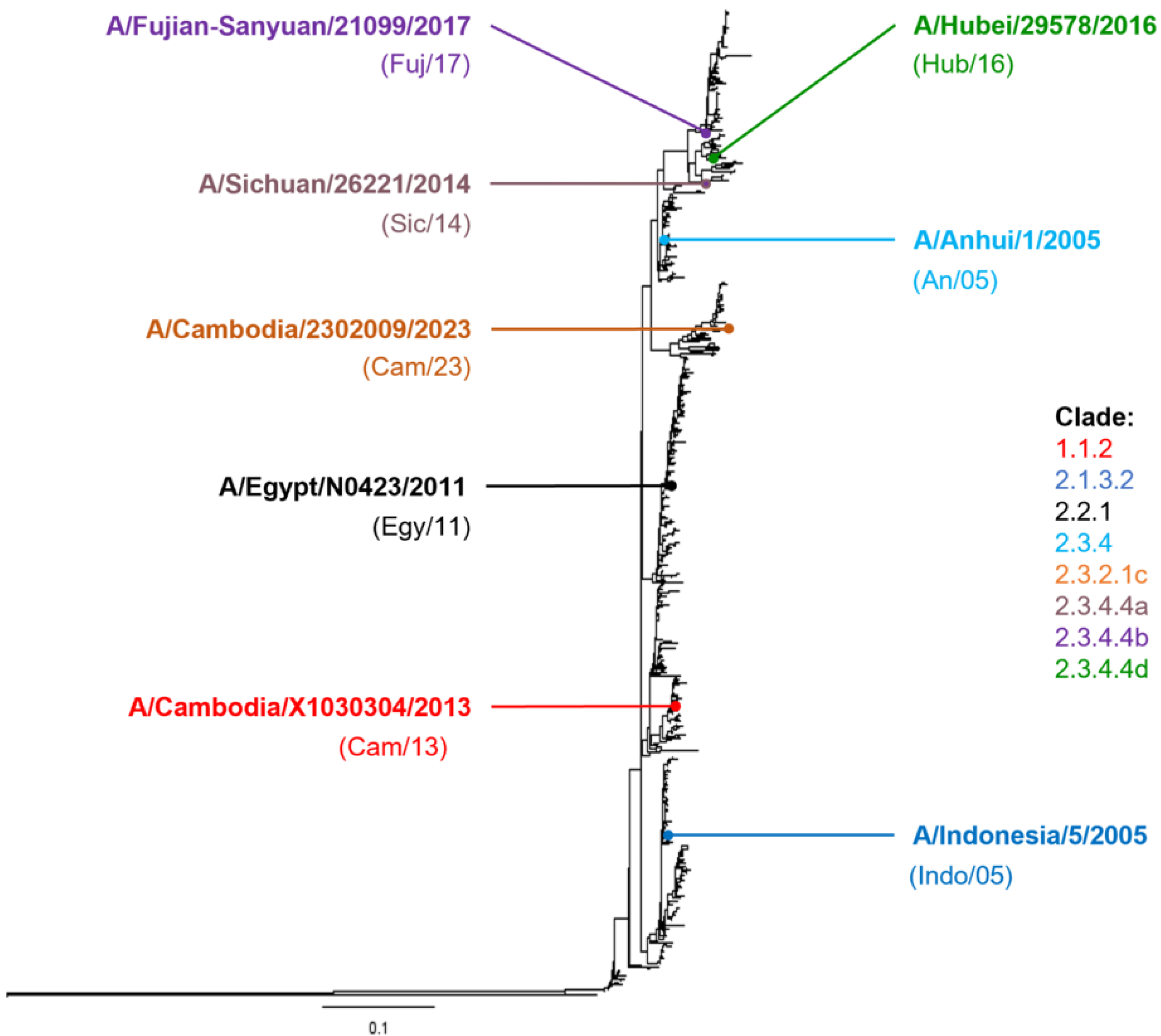
Phylogenetic tree of H5Nx human isolates spanning 2003-2023. Viral strains selected for this study are indicated and colour coded based on their phylogenetic clade.

**Figure 2.**
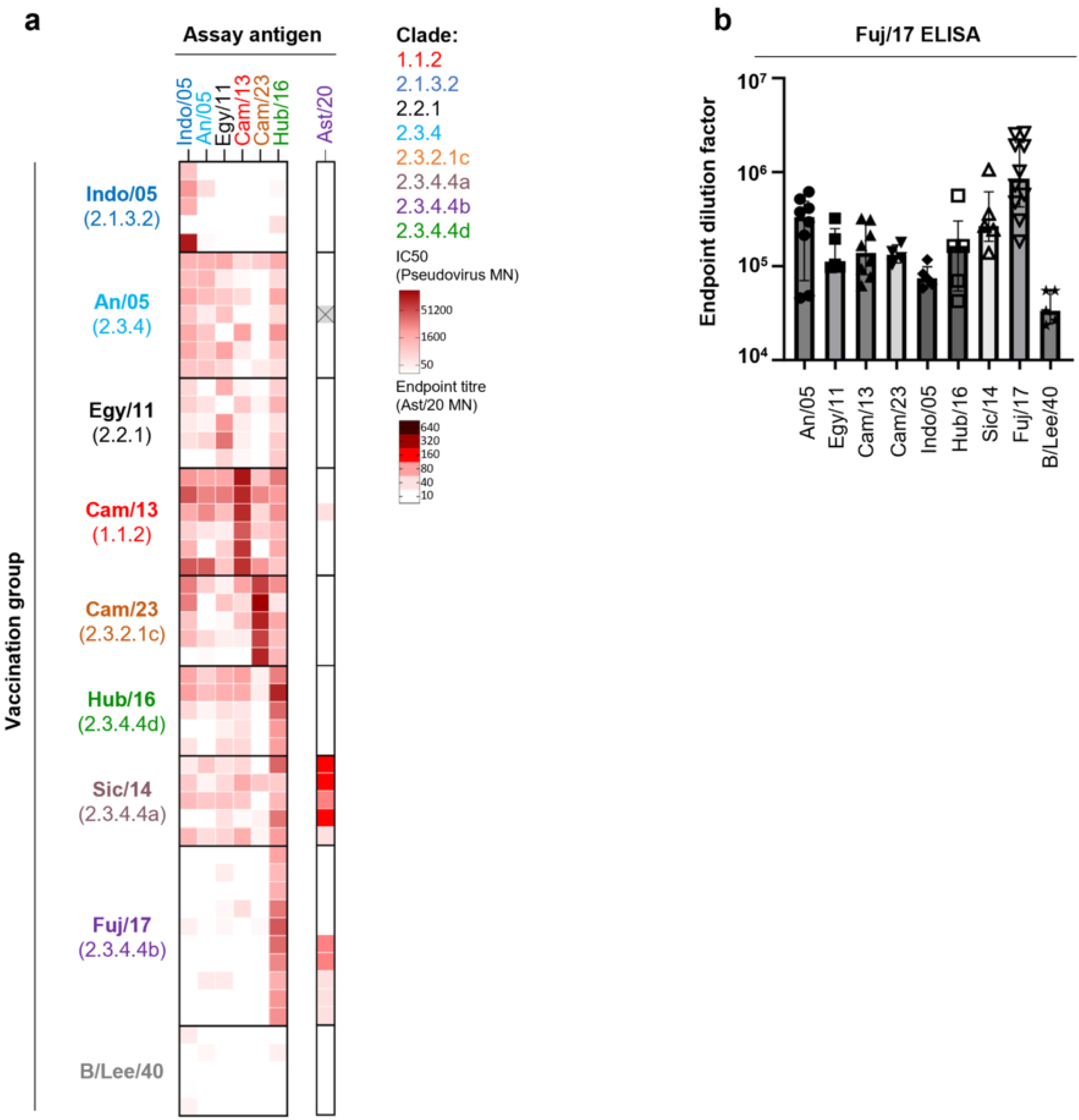
Heterogeneity in cross-strain neutralisation induced by different H5 strains. **(a)** C57BL/6 mice (n=5–10 per group) received two doses of H5 formulated with AddaVax, and antisera were collected 14 days after boosting. Neutralisation activity was assessed using micro-neutralisation (MN) assays with virus pseudotypes (An/05, Egy/11, Cam/13, Cam/23, Indo/05, Hub/16) and shown as serum dilution factor to 50% viral inhibition (IC50). Neutralisation activity against the A/Astrakhan/3212/2020 (H5N8-PR8) virus was examined by MN assays and shown as endpoint neutralising titre. **(b)** Binding reactivity of each H5Nx antisera against Fuj17 full-length HA was measured by ELISA, with data shown as median ± interquartile range.

Marked heterogeneity in cross-strain neutralisation was observed between H5 vaccine strains (Fig. 2a). Immunisation with the Cam/13 or Cam/23 HA elicited broad neutralisation activity against most pseudotypes tested, but not against the Ast/20 live virus. Conversely, immunisation with Indo/05 elicited a highly focussed response largely limited to the vaccinating strain, with An/05, Egy/11 and Hub/16 immunogens driving intermediate cross-strain neutralisation. No cross-neutralisation against the 2.3.4.4b Ast/20 was detected by any antisera, other than the antigenically adjacent 2.3.4.4 antigens (Sic/14 and Fuj/17), highlighting the significant antigenic barrier between H5 clusters. Vaccination with 2.3.4.4 immunogens further highlighted subclade-specific differences in the ability to induce neutralisation breadth, with the 2.3.4.4a Sic/14 immunogen eliciting broad cross-neutralisation, while Fuj/17 from 2.3.4.4b only cross-neutralised the 2.3.4.4d Hub/16 but not non-2.3.4.4b pseudoviruses. Observed differences in cross-neutralisation breadth were not a function of gross immunogenicity, with most HA immunogens eliciting comparable neutralising responses against their cognate pseudovirus (Supp. Fig. 1; Supp. Fig. 2), and with the elicitation of binding antibody responses in all vaccinated serum with some recognition against Fuj/17 (Fig. 2b).

### Impact of vaccine composition on breadth and potency of H5Nx cross-neutralisation

We next investigated vaccine strategies that may broaden antibody cross-neutralisation of H5Nx. Specifically, we examined either vaccination with a stabilised HA mini-stem construct derived from A/Vietnam/1203/2004 (H5-Stem) (22), or polyvalent formulations of full-length H5 HA immunogens as an admix of three (Indo/05, Cam/23, Fuj/17; 3x admix) or eight H5 HAs plus the H5-stem (Indo/05, An/05, Egy/11, Cam/13, Cam/23, Sic/14, Hub/16, Fuj/17, Viet/04 stem; 9x admix) (Fig. 3a).

**Figure 3.**
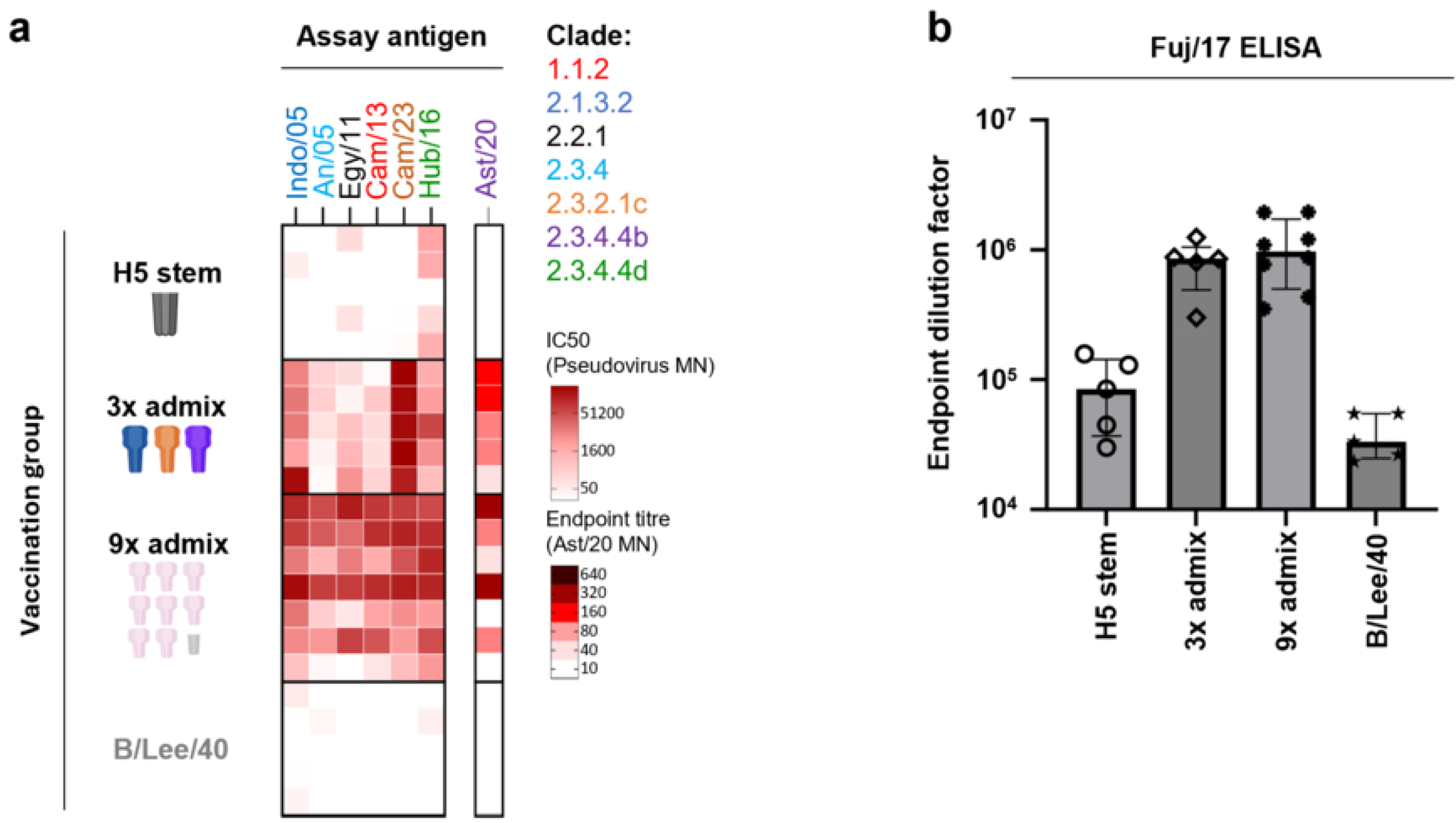
Multivalent H5Nx vaccine formulations enhance the breadth and potency of cross-strain neutralisation. **(a)** C57BL/6 mice were prime-boost immunised with either a Viet/04 HA stem immunogen or admix formulations of full-length HA (3x admix, 9x admix). Cross-strain neutralisation against H5 pseudotypes and A/Astrakhan/2020 was measured; each row represents individual mouse antiserum. **(b)** Binding antibody titres against 2.3.4.4b Fuj/17 HA were measured using ELISA, with data shown as median ± interquartile range.

Vaccination with the stabilised H5 stem did not raise significant neutralising responses against the Ast/20 virus or any of the pseudoviruses tested, apart from Hub/16 which appears uniquely susceptible to antibody-mediated neutralisation. The maximalist 9x admix vaccine drove broad and potent cross-neutralisation against all viruses in our panel, despite the reduced per strain dose (approximately one-tenth) compared to animals receiving a single HA immunogen. This dose-sparing effect was also evident in strain-specific ELISA titres against Fuj/17, where the 9x admix and 3x admix groups showed comparable levels of anti-Fuj/17 antibodies (Fig. 3b). Reduction of vaccine components to three strains maintained neutralising breadth, albeit with a minor reduction in potency against non-cognate strains (Fig. 3a; Supp. Fig. 1; Supp. Fig. 2).

Overall, we found that rational selection of a limited admix of HA immunogens is a tractable pathway to eliciting neutralisation breadth against H5 subtypes, although the strain-specificity of each HA in eliciting antibody breadth remains a selection challenge.

### Ferritin nanoparticles boost H5Nx cross-neutralisation responses

Nanoparticle-based vaccine platforms have been widely reported as an effective strategy to enhance the immunogenicity of subunit influenza vaccines (22–26). To assess whether nanoparticle display could improve the breadth or potency of H5 HA-induced antibody responses, we engineered self-assembling protein nanoparticles as previously described (25) by fusing Cam/23 or Fuj/17 HA to *Helicobacter pylori* ferritin (Cam/23 Ferritin-NP and Fuj/17 Ferritin-NP).

In line with the neutralisation breadth induced by soluble HA immunogens (Fig. 2), we observed broader serum neutralisation activity elicited by Cam/23 Ferritin-NP, and narrower, near clade-specific activity elicited by Fuj/17 Ferritin-NP (Fig. 4a; Supp. Fig. 1; Supp. Fig. 2). Although an admix of both nanoparticles (Ferritin-NP admix) displayed limited capability to further enhance the potency of the response, more robust neutralisation against Ast/20 was found across the three nanoparticle formulations compared to Fuj/17 soluble protein vaccination (Fig. 2a).

**Figure 4.**
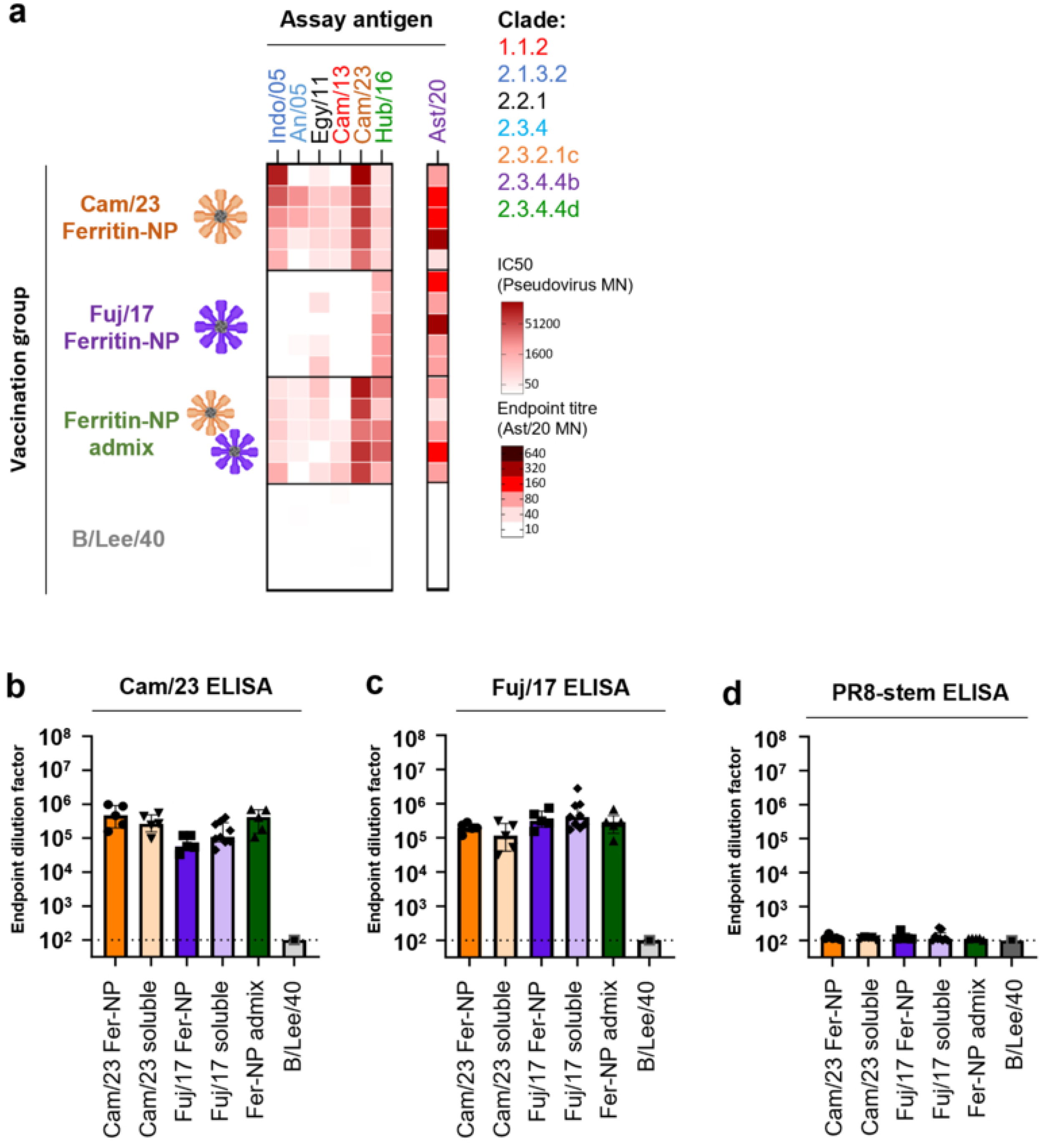
Display on ferritin nanoparticle vaccine platforms can increase the potency of H5Nx cross-strain neutralisation. **(a)** C57BL/6 mice were immunised with ferritin nanoparticle HA vaccines (Cam/23 Ferritin-NP, Fuj/17 Ferritin-NP or a Ferritin-NP admix). Neutralisation activity against H5 pseudotypes and A/Astrakhan/2020 was assessed by MN assays. Binding serum antibody titres against **(b)** Cam/23 or **(c)** Fuj/17 full-length HA were assessed by ELISA. (d) HA stem-specific antibody titres were evaluated against a trimeric PR8 stem protein construct by ELISA. All ELISA assays are graphed by endpoint dilution factor, with data shown as median ± interquartile range.

Increased neutralisation activity driven by nanoparticle display did not appear to be a function of gross immunogenicity, with comparable high titre serum binding responses observed between Ferritin-NP and soluble HA immunised mice (Fig. 4b-4c; Supp. Fig. 3). Similarly, neutralisation breadth was not due to increased antibody recognition of the HA stem, with neither soluble HA protein nor HA ferritin nanoparticle vaccines eliciting robust responses against a conserved stabilised stem domain from the H1N1 strain A/PR8/8/1934 (Fig. 4d). Therefore, nanoparticle display appears to qualitatively enhance the neutralising proportion of the serological response to H5 vaccine.

### Prior immunisation but not infection boosts cross-clade neutralising antibody responses elicited by an H5 Fuj/17 HA vaccine

Human populations have extensive pre-existing immunity to influenza from prior infection and/or immunisation. We therefore investigated the impact of prior immunity upon responses to H5 vaccination. In mice subject to infection with either A/PR8 (H1N1), A/X31 (H3N2) or B/Aus/21 viruses prior to prime-boost vaccination with Fuj/17 HA (Fig. 5a), we saw limited to no impact on serological cross-neutralising activity relative to mock-infected PBS controls (Fig. 5b). In terms of gross immunogenicity, we saw limited impact of infection upon Fuj/17 vaccine-induced serum antibodies recognising Fuj/17 (Fig. 5c) or a heterologous Cam/23 HA (Fig. 5d) before and after receiving the follow-up Fuj/17 vaccination, despite a slight elevation of Fuj/17 HA recognition in the PR8 pre-vaccinated mice (Fig. 5c).

**Figure 5.**
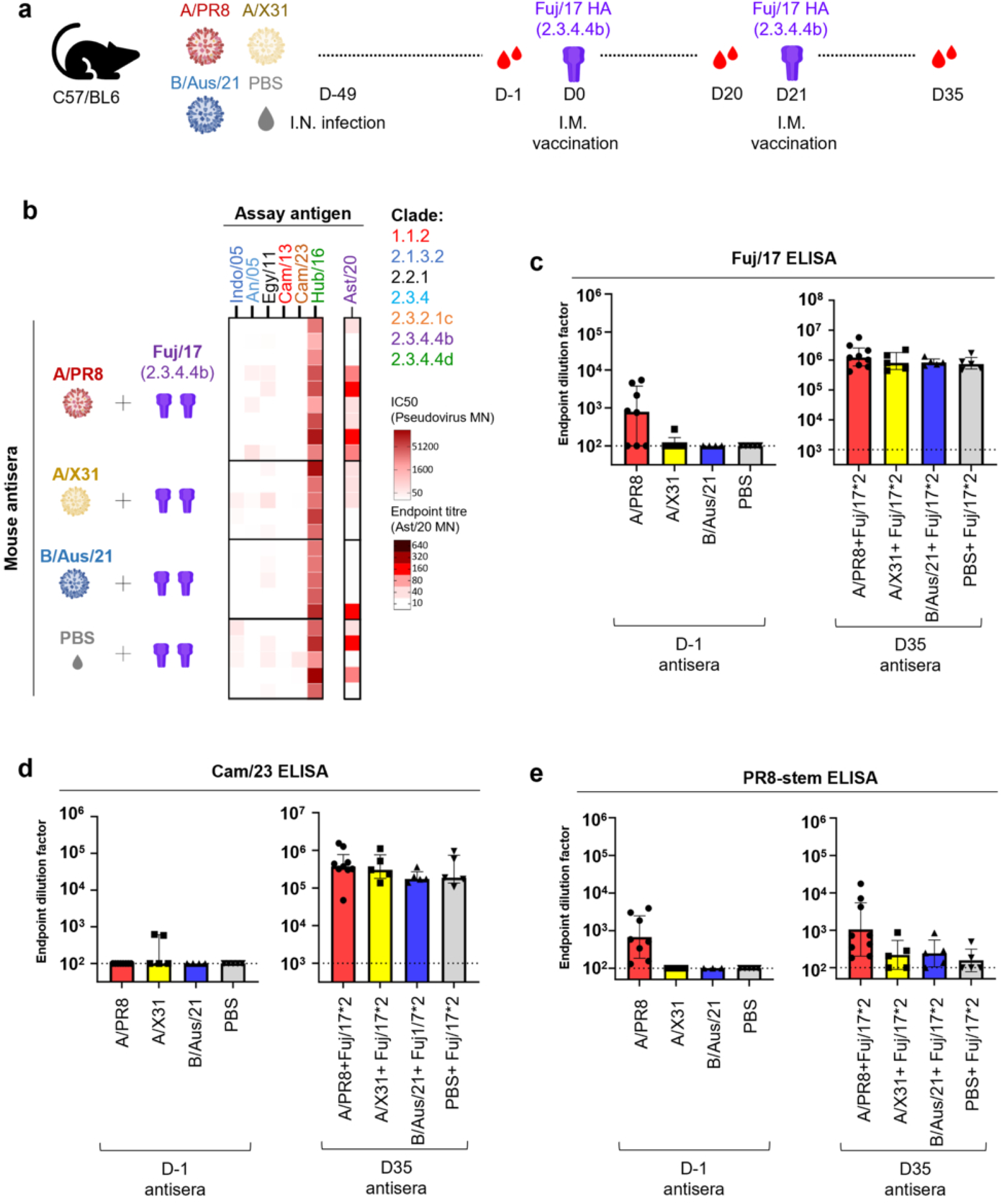
Pre-infection with seasonal influenza strains does not impact immunogenicity of a 2.3.3.4b prime-boost vaccine. **(a)** C57BL6 mice (n=5–10 each group) were infected intranasally with either PR8 (H1N1), X31 (H3N2), or B/Aus21 (B/Austria/2021) 49 days prior to prime-boost intramuscular immunisation with Fuj17 HA in AddaVax. **(b)** Cross-neutralisation against H5Nx viruses was examined by MN assays. Serological binding antibody titres were measured by ELISA for **(c)** Fuj/17, **(d)** Cam/23, or **(e)** the H1N1 stabilised stem domain from A/PR8 (H1N1). ELISA data were shown as median ± interquartile range and compared using Mann–Whitney U tests.

Antibody cross-reactivity within Group 1 and Group 2 IAV viruses has been widely described and is largely concentrated within the conserved HA stem domain (27–29). While we observed a modest increase in H1-stem-specific antibodies in mice infected with the H1N1 A/PR8 virus, this was not robustly boosted by prime-boost Fuj/17 HA vaccination (Fig. 5e).

We next examined the impact of immune memory established by seasonal influenza vaccines on subsequent H5 immunisation. Mice were immunised twice with Southern Hemisphere quadrivalent influenza vaccines (QIV) manufactured for the 2024 [QIV(2024)] or 2025 [QIV(2025)] flu seasons prior to receiving two doses of Fuj/17 HA formulated in Addavax (Fig. 6a). In control mice receiving four doses of QIV(2025), some mice developed modest titres of H5 neutralisation limited to non-2.3.4.4b pseudoviruses (Fig. 6b). In contrast, animals previously vaccinated with either 2024 or 2025 QIV formulations developed broad cross-reactive H5 neutralisation upon immunisation with H5 Fuj/17. In terms of gross immunogenicity, all mice receiving Fuj/17 HA developed robust serum titres of antibody against homologous Fuj/17 (Fig. 6c) or heterologous Cam/23 (Fig. 6d) irrespective of prior QIV administration. All mice immunised with QIV mounted robust antibody responses against CA/09 H1N1pdm HA (Fig. 6e), a virus antigenically similar to the component H1N1 in the vaccines. In contrast to H1N1 infection, animals also elicited robust serum antibody responses against the H1 stem (Fig. 6f), responses that were boosted upon receipt of Fuj/17 HA. Overall, these data suggest seasonal QIV may prime stem-specific immune memory that can be recalled by H5 immunisation and contribute to broader cross-neutralisation of H5 strains.

**Figure 6.**
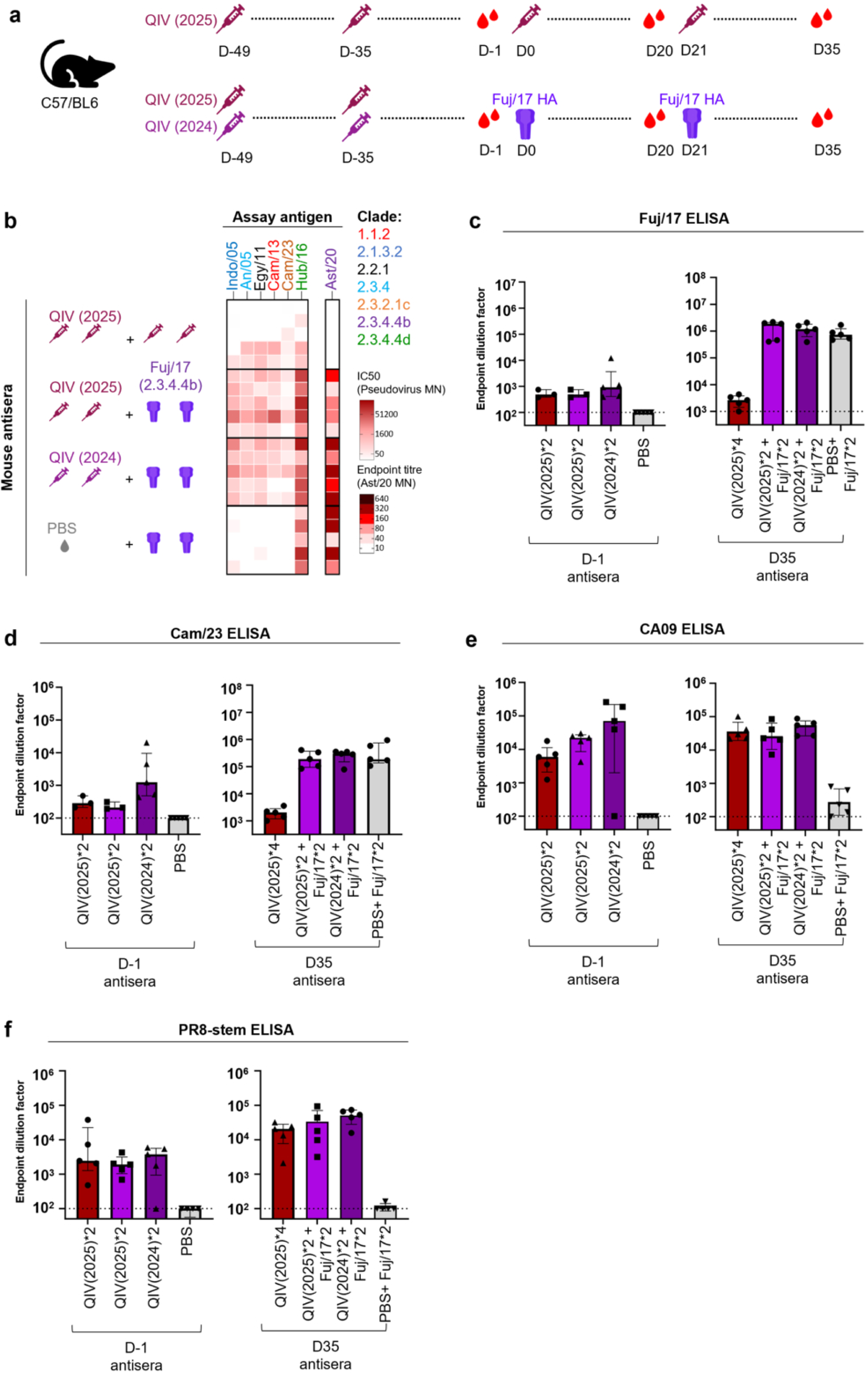
Prior immunisation with seasonal QIV augments the development of cross-strain neutralisation in mice receiving a 2.3.3.4b prime-boost vaccine. **(a)** C57BL/6 mice were immunised twice intramuscularly with seasonal QIV (5ug of total HA content, adjuvanted) prior to receiving prime-boost immunisation with Fuj/17 HA formulated with AddaVax. **(b)** Serum neutralisation activity against H5 was assessed by MN assays. Serological binding antibody titres were measured by ELISA for **(c)** Fuj/17 HA, **(d)** Cam/23 HA, **(e)** H1N1 CA09 HA or **(f)** PR8 stem. ELISA data were shown as median ± interquartile range.

## Discussion

The growing threat of H5 avian influenza, in particular increasing mammal-to-mammal transmission, has created an urgent need for effective and durable H5 vaccines. Current vaccine development faces a critical challenge: designing formulations that provide broad protection against the substantial antigenic and genetic diversity of circulating H5 viruses. Previous studies have extensively mapped such antigenic relations for A(H5) using ferret or guinea pig antisera. Zhang *et al.* resolved 3 putative antigenic clusters, one encompassing ancestral (pre-2.3.4.4) clades, a second cluster primarily comprised of 2.3.4.4 viruses, and a third cluster made of 2.3.4.4.h viruses (30). Kok *et al.* similarly reported 2.3.4.4 and 2.3.4.4b viruses display antigenic differences to the other H5 clades (14). We observed limited cross-neutralisation between pre-2.3.4.4 and 2.3.4.4b viruses in mice vaccinated with different H5 HA immunogens, consistent with the antigenic divergence between these groups. However, we also observed marked strain-specificity in the capacity of different HA immunogens to elicit neutralisation breadth, suggesting predictions of vaccine efficacy cannot be informed solely by antigenic cartography. Asymmetrical cross-neutralisation was driven by discordance between HA antigenicity and immunogenicity, with some viruses broadly neutralised by heterologous antisera, but their corresponding homologous HA antisera failed to induce heterologous neutralising breadth when administered as a putative vaccine immunogen. A Similar phenomenon has been reported by Amanat *et al.* in a SARS-CoV-2 study, in which spike antisera raised against the antigenically central variant P.1 presented broader cross-neutralising activity, while the P.1 virus itself was not efficiently neutralised by heterologous antisera (31). Further clarity is required around the immunological drivers of discrete epitope selection in response to vaccination in order to increase our ability to predict immunogen performance in vivo.

Given the extensive antigenic diversity of H5 viruses with capable of infecting humans and their potential to cause future pandemics, we need strategies to most efficiently elicit broad vaccine coverage. Prior studies have suggested that incorporating specific amino acid mutations (T134A, Q222L and G224S) or removing putative glycosylation sites (T156A) (14) can enhance the antigenic “centrality” of a vaccine immunogen. However, we observed that polyvalent HA admixtures significantly outperformed any given monovalent vaccine in the elicitation of breadth, with both 3x and 9x admix formulations significantly enhancing neutralisation capacity against all representative strains. While defining a minimal admix to provide analogous coverage in humans may require extensive clinical validation, such an approach may bypass the unpredictable immunogenicity of any single individual strain and derisk the utility of stockpiled H5 vaccines in the event of an outbreak.

The conserved HA stem domain presents an attractive target for developing broadly reactive influenza vaccines, as stem-specific antibodies have been reported to be capable of mediating cross-protection against multiple viral subtypes or groups by interfering with viral fusion and/or promoting viral clearance (22, 32). However the HA stem exhibits an intrinsic immunological subdominance and is typically outcompeted by immunodominant responses directed against the variable HA head (33). This limitation is evidenced by the low levels of stem-targeting antibodies observed in recipients of inactivated H5N1 vaccines (34, 35). These challenges have driven the development of advanced stem-based immunogen designs, which often shield or remove the hypervariable head domain to restore robust stem-specific antibody generation and confer *in vivo* protection against diverse viral challenges (36–41). In the current study, we saw limited elicitation of cross-neutralising antibody responses boosted by a prototypic stabilised H5 stem immunogen based on a design established for H1N1 (22), suggesting that this immunogen may poorly recapitulate the antigenic structure of the H5 stem. Notably, we did see that prior immunity established through seasonal influenza vaccination elicited robust binding titres against the H1 stem from PR8, and that these stem responses were boostable by subsequent H5 Fuj/17 immunisation and likely contributing to the broader cross-neutralisation observed. Previous studies have also demonstrated that seasonal influenza vaccination can facilitate protective cross-subtype antibody responses following H5 vaccination in various pre-clinical models (42–44). Interestingly, we did not see analogous enhancement of neutralisation breadth in animals subject to prior infection, including with an H1N1 strain. Together, these findings suggest that the antigenicity of the Group 1 stem domain is complicated and can dramatically influence the ability to focus antibody responses onto rare epitopes able to mediate cross-neutralisation of H1 and H5 viruses.

In summary, we find that rational strain selection, minimal multiplexing, nanoparticle display and exploitation of HA stem-directed immune memory are all factors that could be exploited to expand the protective breadth of prototypic HA-based vaccines against avian H5 viruses.

## Materials and Methods

### Protein generation

Recombinant influenza A virus hemagglutinin proteins were produced as previously described (21). Full-length HA ectodomains were engineered with C-terminal fusion tags comprising a trimerization motif from bacteriophage T4 fibritin foldon domain, the biotinylation-competent AviTag peptide sequence GLNDIFEAQKIEWHE, and a histidine purification tag (complete sequences provided in Supplementary Data_H5 protein sequence). Codon-optimized genes encoding IAV HA ectodomains [A/Indonesia/5/2005 (EPI_ISL_5729), A/Anhui/1/2005 (EPI_ISL_10058), A/Egypt/N0423/2011 (EPI_ISL_120281), A/Cambodia/X1030304/2013 (EPI_ISL_153029), A/Cambodia/2302009/2023 (EPI_ISL_17069010), A/Sichuan/26221/2014 (EPI_ISL_163493), A/Fujian-Sanyuan/21099/2017 (EPI_ISL_304404), and A/Hubei/29578/2016 (EPI_ISL_341293)] were incorporated a Y–F substitution (at positions 95-98 depending on the strain) and a furin cleavage site, synthesised commercially (GeneArt) and cloned into mammalian expression plasmids. Stabilised stem constructs derived from A/VietNam/1203/2004, A/California/07/2009 and A/Puerto Rico/8/1934 (PR8) hemagglutinin were generated following design principles from prior studies (22). All recombinant proteins were produced via transient transfection of Expi293 suspension cells (Life Technologies, A14527) and subsequently purified through 6×histidine-tag affinity chromatography, followed by size-exclusion chromatography. Protein sizes were validated on SDS-PAGE chromatography (Supp. Fig. 4).

### Generation of Ferritin HA nanoparticles

Recombinant HA-ferritin nanoparticles were generated following previously established protocols (23). Briefly, HA ectodomains from A/Fujian-Sanyuan/21099/2017 and A/Cambodia/2302009/2023 were engineered to incorporate a Y–F substitution (at positions 95-98 depending on the strain) and ablation of the furin cleavage site. These modified HA sequences were fused to the C-terminus of a gene encoding Helicobacter pylori non-haem iron-containing ferritin (GenBank accession NP_223316) via a Gly-Ser-Gly tripeptide linker and subcloned into mammalian expression vectors. Sequence-verified plasmids were transfected into Expi293F suspension cells (Life Technologies, Thermo Fisher Scientific) according to the manufacturer’s protocol. Culture supernatants were harvested 4–5 days post-transfection, centrifuged, and subjected to anion exchange chromatography using a HiTrap Q HP column (Cytiva). Nanoparticle preparations underwent further purification by size-exclusion chromatography on a HiScreen Capto^®^ Core 400 column (Cytiva). Expression was confirmed by SDS-PAGE.

### Intramuscular immunisation of mice with recombinant proteins

All animal work was conducted in accordance with guidelines set by the University of Melbourne Animal Ethics Committee (ethics approval number 21799). Mouse antisera were generated by intramuscular immunisations of C57BL/6 mice with recombinant hemagglutinin antigens (full-length HA, HA stem domain, or HA-ferritin nanoparticles). Each vaccine dose contained 5 μg of purified protein in phosphate-buffered saline (PBS; final volume 50 μL) emulsified 1:1 with Addavax (InvivoGen) (total injection volume 100 μL). For multivalent formulations, equal amounts of each HA were combined to a total protein content of 5 μg per dose. Animals received a booster dose at week 3 following primary vaccination, with serum collected on day 14 post-boost.

### Intramuscular immunisation of mice with seasonal influenza vaccines

Pre-immunity was established by seasonal influenza vaccination using commercially available QIV (Flucelvax^®^ Quad, Seqirus). Two seasonal formulations were employed: the 2024 Southern Hemisphere vaccine containing A/Wisconsin/67/2022 (H1N1)pdm09-like, A/Massachusetts/18/2022 (H3N2)-like, B/Austria/1359417/2021 (Victoria lineage)-like, and B/Phuket/3073/2013 (Yamagata lineage)-like virus antigens; and the 2025 Southern Hemisphere vaccine containing A/Wisconsin/67/2022 (H1N1)pdm09-like, A/District of Columbia/27/2023 (H3N2)-like, B/Austria/1359417/2021-like, and B/Phuket/3073/2013-like virus antigens. QIV prime-boost regimens consisted of 5 μg total HA content per dose in an equal volume of Addavax (InvivoGen), administered at a two-week interval. Fuj/17 H5 immunisation, as described above, was subsequently given five weeks following the QIV boost. Serum samples were obtained at three timepoints: day -1 (24 hours prior to Fuj/17 H5 priming), day 20 (24 hours prior to Fuj/17 H5 boost), and day 35 (study endpoint).

### Intranasal infection of mice

To establish pre-existing immunity through viral infection, C57BL/6 mice were challenged intranasally with either A/Puerto Rico/8/1934 (H1N1; 20 PFU), A/Aichi/2/1968 x A/Puerto Rico/8/1934 reassortant (X-31, H3N2; 1,000 PFU), or B/Austria/1359417/2021 (IBV; 1×10⁵ PFU). Viruses were administered in 50μL of PBS under isoflurane anaesthesia. At 7 weeks post-challenge, mice received Fuj/17 H5 immunisation and sera were collected as described above.

### Generation and titration of H5 pseudotypes with lentiviral vectors

H5-pseudotyped lentiviral particles (PVs) were generated as previously described(45). HA ectodomains from A/Indonesia/5/2005, A/Anhui/1/2005, A/Egypt/N0423/2011, A/Cambodia/X1030304/2013, A/Cambodia/2302009/2023, and A/Hubei/29578/2016 were fused to the transmembrane domain of A/Indonesia/5/2005 (sequence provided in Supplementary Data_H5 protein sequence) and cloned into pCMV expression plasmids. To facilitate viral release, a plasmid expressing a soluble version of the neuraminidase (NA) ectodomain from A/Indiana/10/2011 (H3N2) was included. Human airway trypsin-like protease (HAT), cloned into pCAGGS, was included to enable HA cleavage during pseudovirus production (46). Lentiviral packaging plasmids encoding gag/pol, tat, and rev were cloned into pHDM or pRC-CMV vectors. A pHAGE2-CMV reporter plasmid co-expressing firefly luciferase and ZsGreen was used for PV production.

HEK293T cells were co-transfected in 6-well plates using Lipofectamine™ 3000 (Thermo Fisher Scientific) with lentiviral packaging plasmids (0.28 μg each), reporter plasmid (1.28 μg), HA expression plasmid (0.44 μg), soluble NA plasmid (0.22 μg), and HAT plasmid (0.275 μg). Transfections were performed in 500 μL reduced-serum Dulbecco’s Modified Eagle Medium (DMEM, Gibco) supplemented with 2% fetal calf serum (FCS) and 1% penicillin/streptomycin/L-glutamine (PSG; Thermo Fisher Scientific) for 6 hours at 37°C with 5% CO₂, followed by replacement with complete DMEM containing 10% FCS and 1% PSG. Supernatants were harvested 72 h post-transfection, filtered through 0.45 μm membranes, and stored at -80°C in single-use aliquots.

Pseudovirus titres were determined by infecting HEK293T cells (1.5 × 10⁴ cells/well) in white 96-well plates with three-fold serial dilutions of PVs (40 μL/well) and incubating for 48 hours at 37°C with 5% CO₂. Firefly luciferase activity was measured using the BriteLite™ Plus assay (PerkinElmer), and PV input was normalised to achieve 100,000 relative luciferase units (RLU).

### Pseudovirus micro-neutralisation assays

Neutralisation activity of murine antisera was tested against H5 pseudovirions. Antisera were diluted 1:50, then serially diluted two-fold in 96-well white polystyrene plates. Each plate included negative control wells (cells only) and positive control wells (cells with PV). Pseudoviruses were diluted to achieve 100,000 RLU in positive control wells, added at 40 μL per well, and incubated with diluted sera for 1 hour at 37°C with 5% CO₂. Subsequently, 40 μL of HEK293T cells (1.5×10⁴ cells per well) were added to each well, and plates were incubated for 48 hours at 37°C with 5% CO₂. Firefly luciferase activity was measured using the BriteLite™ Plus assay (PerkinElmer). Neutralisation activity was determined as the half-maximal effective dilution factor (IC₅₀) required to inhibit luciferase expression by 50% relative to positive control wells.

### Micro-neutralisation assay

A recombinant IAV with the HA and NA from A/Astrakhan/3212/2020 (clade 2.3.4.4b H5N8) with the remaining gene segments derived from A/Puerto Rico/8/1934 was used for live virus micro-neutralisation as previously described (47). Madin-Darby Canine Kidney (MDCK) cells (ATCC CCL-34) were plated at a density of 5×10⁴ cells per well in 96-well tissue culture plates and incubated overnight at 37°C in complete growth medium consisting of DMEM supplemented with 10% FCS, 1% PSG, and 1% sodium pyruvate (Gibco). Mouse serum samples were heat inactivated at 56°C for 30 minutes, and then serially diluted two-fold (1:10–1:1280) in virus infection medium [VIM; DMEM containing 1% PSG, 1% sodium pyruvate, and 0.1% L-1-tosylamido-2-phenylethyl chloromethyl ketone (TPCK)-treated trypsin]. Diluted sera were incubated with 200 TCID₅₀ of the A/Astrakhan/3212/2020 virus for 60 minutes at 37°C. After incubation, MDCK cells were washed twice in PBS. VIM (100 μL) supplemented with 2 μg/mL TPCK-treated trypsin and 100 μL virus–serum mix were added to the MDCK monolayer. Control wells of virus alone and VIM alone were included on each plate. Virus input was back titrated on MDCK cells. Cells were incubated for 72 hours at 37°C with 5% CO₂. The presence of virus was determined by haemagglutination assay. Briefly, 25 μL of supernatant from each well was mixed with 25 μL of 1% (v/v) turkey erythrocytes and incubated for 30 min at room temperature, and the presence of virus was recorded. Micro-neutralisation titres were determined as the reciprocal of the highest dilution at which virus neutralisation (negative in HA assay) was observed.

### ELISA (enzyme-linked immunosorbent assay)

Antibody binding to HA or stabilised stem proteins was assessed by capture ELISA. Briefly, high-binding 96-well microplates (Thermo Fisher Scientific) were coated overnight at 4°C with anti-His-tag polyclonal antibody (GenScript) at 2 μg/mL in PBS. Wells were blocked with 5% (w/v) skim milk powder in PBS for 1 hour at room temperature, followed by three washes with PBS containing 0.05% Tween-20 (PBST). Recombinant HA proteins diluted to 2 μg/mL in blocking buffer were added to wells and incubated for 1 hour at room temperature. Following washing, serially diluted murine antisera were added and incubated for 2 hours at room temperature. After washing, horseradish peroxidase (HRP)-conjugated anti-mouse IgG secondary antibody (KPL) diluted 1:20,000 in blocking buffer was added and incubated for 1 hour at room temperature. Plates were washed extensively with PBST, and bound antibody was detected using 3,3’,5,5’-tetramethylbenzidine (TMB) substrate solution (Sigma-Aldrich). Reactions were stopped with 1 M sulphuric acid, and absorbance was measured at 450 nm using a microplate reader. HA-specific antibody binding was quantified by calculating the endpoint dilution giving a signal 2×background using a fitted curve (four-parameter logistic regression).

### Statistical Analyses

Data is presented as median and interquartile range, graphed using Prism ver 5.0 (GraphPad).

## Authors’ contributions

YCL, SJK, MK and AKW designed the study. YCL, AK, RE, WSL performed experiments. YCL and MK performed serological analyses. YCL, MK and AKW analysed data. YCL, AKW and MK wrote the manuscript. All authors read and revised the manuscript.

## Acknowledgements

The work has been generously supported by the Australian National Health and Medical Research Council. YCL is supported by the Melbourne graduate research scholarship, provided by University of Melbourne. We are grateful to the WHO Collaborating Centre for Reference and Research on Influenza in Melbourne for providing the A/Astrakhan/3212/2020-PR8 virus used in neutralisation assays. We thank Carol Weiss for providing the plasmid encoding HAT. The funders had no role in study design, data collection and analysis, decision to publish or preparation of the paper. For the purposes of open access, the author has applied a CC BY public copyright licence to any Author Accepted Manuscript version arising from this submission.

## Figure legends

**Supp. Fig. 1. Pseudovirus MN titres of mouse antisera immunised with H5 HA.** Data are shown as median ± interquartile range.

**Supp. Fig. 2. Ast/20 MN titre of mouse antisera immunised with H5 HAs.** Data are shown as median ± interquartile range.

**Supp. Fig. 3. ELISA titer of mouse antisera immunised with HA Ferritin-NP against H5 HAs and PR8 stem.** Data were shown as median ± interquartile range.

**Supp. Fig. 4. SDS-PAGE of expressed recombinant soluble H5 proteins**. Lane 1: marker, lane 2: 3µg A/Anhui/1/2005 HA trimer, lane 3: 3µg A/Cambodia/2302009/2023 HA trimer, lane 4: 3µg A/Egypt/N0423/2011 HA trimer, lane 5: 3µg A/Fujian-Sanyuan/21099/2017 HA trimer, lane 6: A/Hubei/29578/2016 HA trimer, lane 7: A/Sichuan/26221/2014 HA trimer, lane 8: A/Cambodia/X1030304/2013 HA trimer, lane 9: marker, lane 10: A/Indonesia/5/2005 HA trimer, lane 11: A/Viet_Nam/1203/2004 HA stem.

